# Disruption of the FGFR1-FGF23-Phosphate Axis and Targeted Therapy in a Murine Model of Osteoglophonic Dysplasia

**DOI:** 10.1101/2025.11.14.680268

**Authors:** Giuliana Ascone, Rajdeep Kaur, Arwaa Mehran, Cecilia Rivas, Rebeca Galisteo, Irene Ginty, Shanna Cloud, Arthur MacLarty, Li Li, Gene Elliot, Mara Riminucci, Alessandro Corsi, Dawn Watkins-Chow, Lisa Garrett, Iris Hartley, Luis Fernandez de Castro, Carlos R. Ferreira

**Author notes:** Corresponding author: Carlos R. Ferreira, 10 Center Drive, Building 10, Room 2-5142, Bethesda, MD 20892, USA. These authors share first authorship (Giuliana Ascone, Rajdeep Kaur).

## Abstract

Osteoglophonic Dysplasia (OGD) is an autosomal dominant skeletal dysplasia characterized by impaired bone growth resulting in short stature, severe craniofacial abnormalities, and in some patients FGF23-mediated hypophosphatemia. It is caused by gain-of-function variants in FGFR1, particularly in or near the transmembrane domain of the receptor. We used CRISPR in mice to knock-in the FGFR1 p.N330I variant, chosen based on its association with FGF23 excess. Skeletal phenotyping of this *Fgfr1^+/N330I^* model demonstrated markedly reduced body weight and naso-anal length, shortened long bones, and craniosynostosis, all hallmarks of the human disease. Mutant mice exhibited profound microarchitectural changes in cortical bone and severe disorganization of the growth plate and articular cartilage, driven by decreased cell proliferation and increased apoptosis in skeletal tissues. In addition to osteochondrodysplasia, we noted dramatic increases in plasma FGF23 and hypophosphatemia, driven by upregulated *Fgf23* expression and protein levels in bone, with consequent undermineralization. An in vivo ossicle assay allowed longitudinal evaluation of mineral metabolism. We modulated the signaling pathway by repurposing an inhibitor of the overactive receptor, infigratinib, resulting in partial restoration of naso-anal length in treated mutant mice. This first model of OGD offers insights into the disease pathogenesis and open avenues for targeted therapeutic strategies.

## Introduction

Fibroblast growth factors (FGFs) and fibroblast growth factor receptors (FGFRs) play a vital role in axial and craniofacial skeletal development (1). FGFs bind to one of the four FGFRs to exert their biological functions. FGFRs are tyrosine kinase transmembrane receptors with an extracellular region containing three immunoglobulin (Ig)-like domains (D1, D2 and D3) that are required for binding to FGFs. The transmembrane domain and the adjacent juxta-membrane intracellular domain participate in dimerization, while the tyrosine kinase sub-domains mediate intracellular signaling (1). Binding of FGF ligands to their cognate FGFRs causes the dimerization of the receptor and activation of the tyrosine kinase domains, resulting in a signaling cascade involved in regulating cell proliferation and differentiation processes (2).

The *FGFR* gene family consists of four genes, *FGFR1, FGFR2, FGFR3, FGFR4* that encode their respective proteins. FGFR1 is mainly expressed in the brain and mesenchymal tissues during embryogenesis, as well as in brain, bone, kidney, skin, lung, heart and muscle in the adult life (3). Alternative splicing generates several transcripts of *FGFR1* that give rise to various tissue-specific isoforms showing different binding affinity and specificity to FGFs (2).

FGFR1, in conjunction with the co-receptor Klotho, is the main mediator of the actions of the hormone FGF23. FGF23 is important in phosphate metabolism, acting through FGFR1-Klotho binding at the renal proximal tubule to downregulate phosphate reabsorption, decreasing blood phosphate and increasing urinary phosphate (4). FGF23 also decreases 1,25-dihydroxy-vitamin D (calcitriol) levels, limiting gut absorption of calcium and phosphate (4). Inappropriately high FGF23 signaling and resulting chronic hypophosphatemia are associated with disorders of FGF23 excess such as tumor-Induced osteomalacia (TIO) in adults and X-linked hypophosphatemic rickets (XLH) in children (5, 6), associated with fatigue, muscle weakness, fractures, and pain. FGF23 production is positively regulated by blood phosphate and calcitriol levels. In osteocytes, FGFR1 is also likely involved in the regulation of FGF23 production, using a positive feedback loop whereby increased FGFR1 signaling results in increased FGF23 secretion.

The functional consequences of excess FGFR1 are manifest in osteoglophonic dysplasia (OGD), caused by gain-of-function (GoF) variants in the fibroblast growth factor receptor 1 (*FGFR1*), causing constitutive receptor activation (7, 8). This rare skeletal disorder is characterized by aberrant bone growth resulting in severe short stature and distinctive craniofacial features. Clinical features of OGD include rhizomelia (disproportionate shortening of the proximal limbs), craniosynostosis (premature fusion of the skull bones), prominent forehead, midface hypoplasia, mandibular prognathism, and impaired tooth eruption (9).

To date, three activating variants in *FGFR1* (NM_023110 .3) have been found in more than one family with OGD, including c.929T>A, p.(Asn330Ile) within the Ig-like III domain (D3) (7, 8), and c.1121A>G, p.(Tyr374Cys) (7, 10) and c.1141T>C, p.(Cys381Arg) (7, 8, 11) within the adjacent transmembrane domain. OGD patients harboring the p.(Asn330Ile)/N330I variant have FGF23-mediated hypophosphatemia (7), but the precise mechanism causing hypophosphatemia and FGF23-excess is not fully understood. Thus, patients with OGD may provide valuable insights into the underlying pathways of phosphate metabolism, which could have repercussions beyond mineral metabolism. Indeed, elevated levels of FGF23 have been associated with left ventricular hypertrophy and cardiovascular mortality (12–14), and hyperphosphatemia is associated with arterial calcification, cardiovascular disease and all-cause mortality (15, 16). Hypophosphatemia from cancer immunotherapy is associated with neurotoxicity (17), and abnormal phosphate concentrations are known to influence the risk of a variety of cancers (18).

Understanding the pathobiology of OGD can provide insights into FGFR1, FGF23, and phosphate metabolism. To pursue these investigations, we generated the first *Fgfr1* murine model that recapitulates the clinical features of OGD, allowing elucidation of the molecular mechanisms involved in OGD and enabling the assessment of future potential therapies.

## Results

### Fgfr1^+/N330I^ mice show involvement of the appendicular skeleton

We used CRISPR/Cas9 technology to generate a knock-in mouse model expressing the disease-causing FGFR1 variant, p.N330I (**Figure 1A**), chosen because of its frequency and its association with FGF23 excess and hypophosphatemia in patients. Morphometric measurements revealed that the *Fgfr1^+/N330I^* mice were significantly smaller compared to their wild type (WT) littermates (**Figure 1B**) and displayed a significant reduction of the total body weight (BW) and whole-body length (47% and 69%, respectively) (**Figure 1C**). Other growth parameters such as the naso-anal and tail length were also significantly reduced in the mutant mice (**Figure 1D**). In addition, the long bones of the hindlimbs of *Fgfr1^+/N330I^* mice were markedly abnormal and shorter when compared to WT controls (**Figure 1E**), with femoral length and tibial length in mutant mice reduced by 58% and 55%, respectively (**Figure 1F**). The femurs from mutant mice also lacked several landmarks, including the femoral head and neck, greater trochanter, lesser trochanter, and third trochanter. Mutant mice showed severely impaired gait due to the significant long bone shortening, as well as hindlimb contractures **(Supplemental Video 1)**.

**Figure 1.**
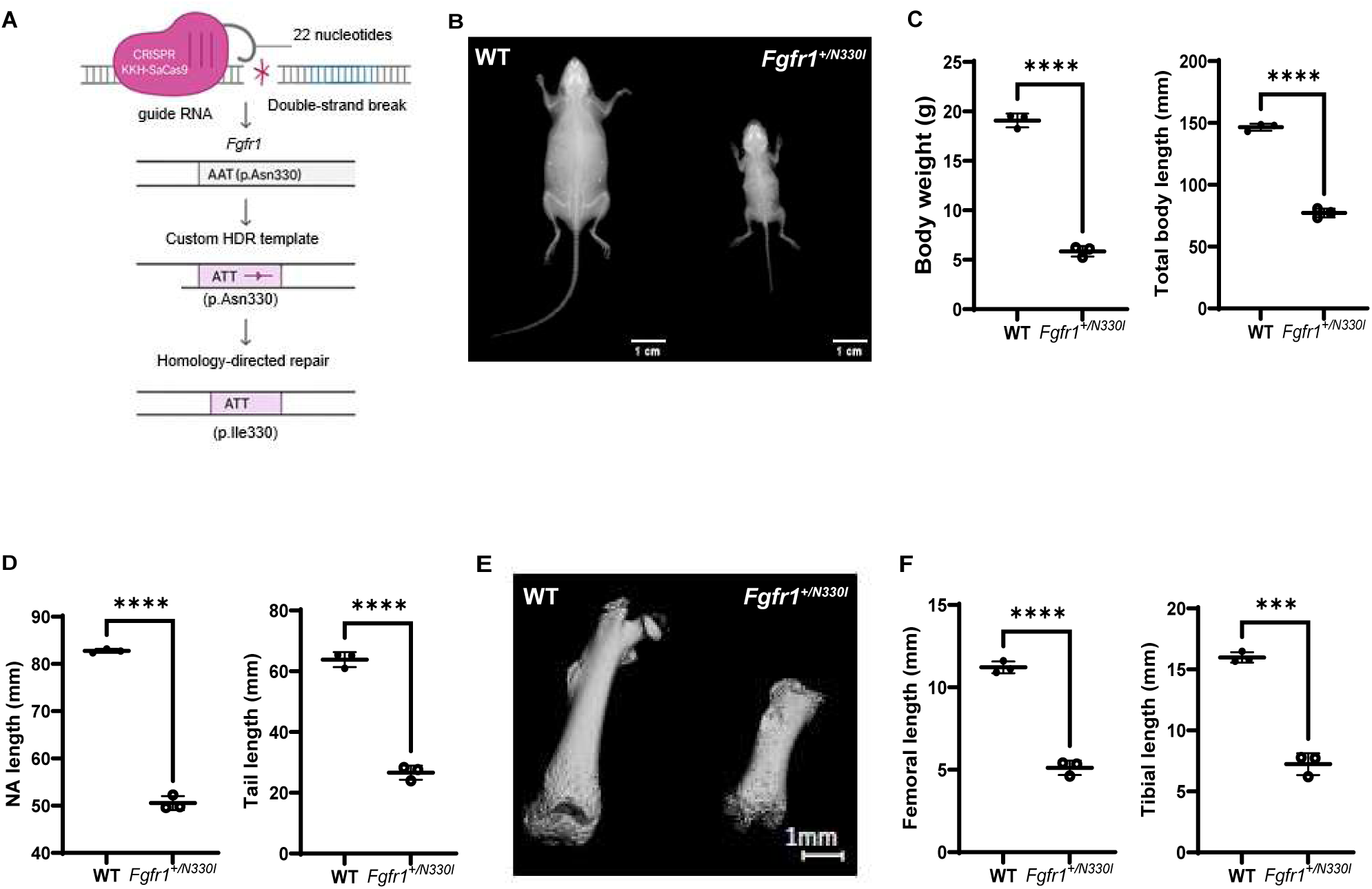
*Fgfr1^+/N330I^* mice show severe growth impairment. **(A)** Design of the knock-in *Fgfr1^+/N330I^* mice. Created with BioRender.com. **(B)** Representative images of whole-body radiography of 5-week-old mice (littermate control on the left; mutant mouse on the right). **(C)** 5-week-old *Fgfr1^+/N330I^* mice are markedly smaller than their WT littermate controls as determined by the reduction of total weight and whole-body length. **(D)** 5-week-old *Fgfr1^+/N330I^* mice have reduced naso-anal and tail length as compared to WT controls. **(E)** Representative microCT scan reconstructed at the same scale, showing severely shortened femur of 3-week-old mutant mouse (right) with lack of typical landmarks (femoral head and neck, greater trochanter, lesser trochanter, third trochanter) as compared to a femur from a littermate control (left). Mutant mice show rhizomelia, or shortening of the proximal appendicular long bones. **(F)** The lengths of both femur and tibia are dramatically reduced in the mutant mice. Horizontal and vertical lines represent the mean ± SD of 3 mice/group (**** p < 0.0001).

We compared the femurs of 5-week-old WT and mutant mice to evaluate differences in bone architecture and microstructure. Ex vivo microCT scans showed lack of trabecular bone in *Fgfr1^+/N330I^* mice and lower cortical bone volume fraction (BV/TV), cortical bone area fraction (Ct.Ar/Tt.Ar), cortex volume, and cortical thickness (Ct.Th) compared to WT controls (**Figure 2A**). Cortical porosity (Ct.Po) was significantly increased in the mutant mice (**Figure 2B**), indicating poor cortical bone microstructure. In line with these findings, *Fgfr1^+/N330I^*mice showed a 13% reduction of femoral bone mineral density (BMD) when compared to their WT littermate controls (**Figure 2C**). This correlated with a reduction of whole-body BMD and whole-body bone mineral content (BMC) (44% and 89% reduction, respectively) (**Figure 2D**). The data indicate that both bone elongation and quality are impaired in the mutant mice.

**Figure 2.**
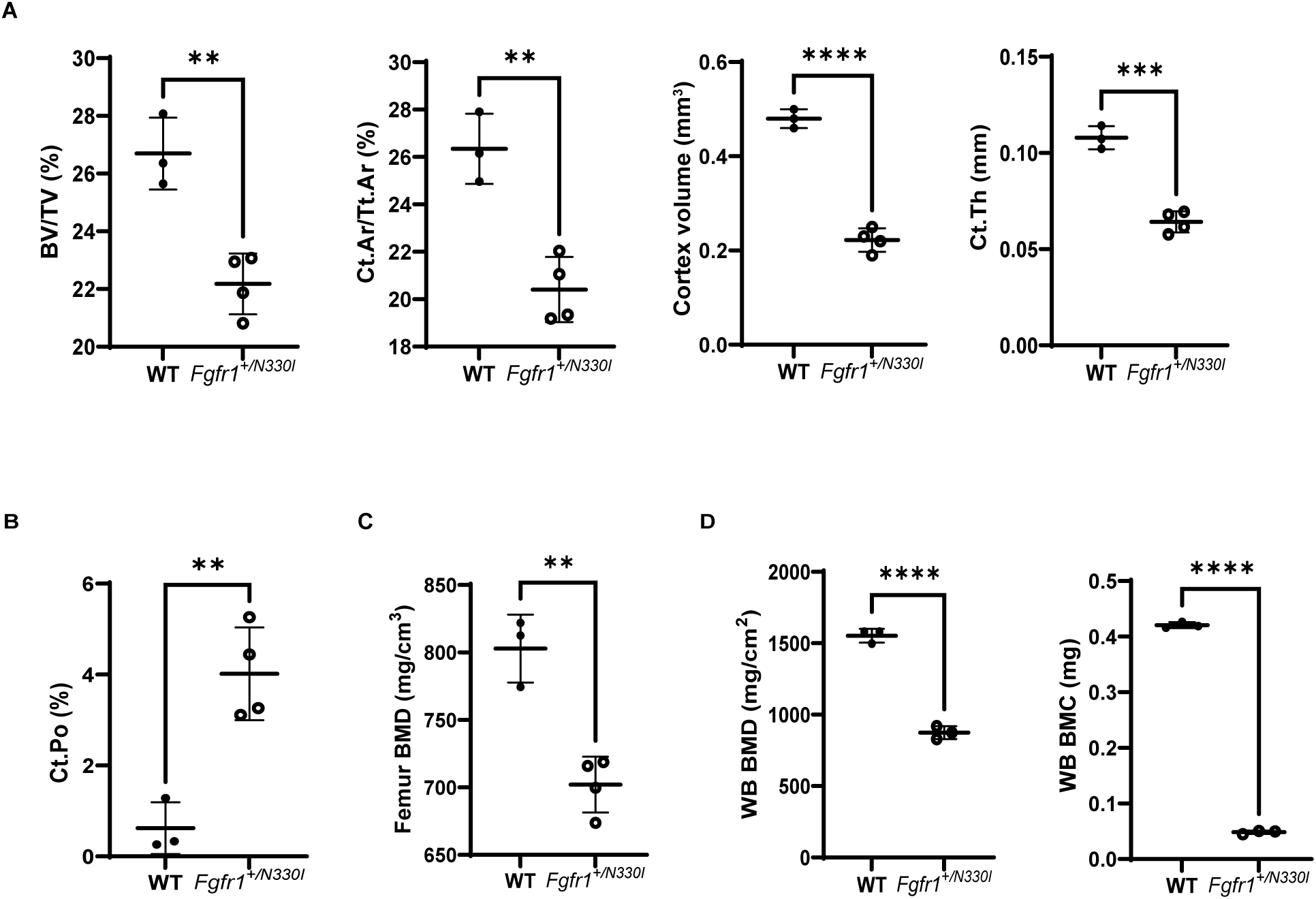
*Fgfr1^+/N330I^* mice show profound alterations of femoral microarchitecture and bone mineral density. **(A)** *Fgfr1^+/N330I^*mice show a significant reduction of bone volume fraction (BV/TV), cortical bone area fraction (Ct.Ar/Tt.Ar), cortex volume, and cortical thickness (Ct.Th) as compared to WT littermate controls (17%, 23%, 54% and 41%, respectively). **(B)** *Fgfr1^+/N330I^* mice show an impaired cortical structure of femurs characterized by an increased cortical porosity (4% vs. 0.6% in WT controls). **(C)** *Fgfr1^+/N330I^* mice display a significant reduction of femur bone mineral density (BMD). Horizontal and vertical lines represent the mean ± SD of 3 WT or 4 mutant mice. **(D)** Whole-body bone mineral density (BMD) and bone mineral content (BMC) are significantly reduced in mutant mice. Horizontal and vertical lines represent the mean ± SD of 3 or 4 mice/group (** p < 0.01, *** p < 0.001; **** p < 0.0001).

### Fgfr1^+/N330I^ mice show involvement of the craniofacial skeleton

Because OGD is accompanied by craniofacial anomalies, we assessed whether the mouse model developed abnormalities in the craniofacial skeleton. In 5-week-old *Fgfr1^+/N330I^* mice skull length was reduced by 35%, eye socket length and width were reduced by 33% and 54%, respectively, and snout length was reduced by 72%, indicating that the pathogenic variant leads to a severe craniofacial phenotype including flat face, similar to that observed in affected patients (**Figure 3A**). At 5 weeks, all sutures were prominently visible in WT mice, whereas *Fgfr1^+/N330I^*mice exhibited premature fusion of frontal and coronal sutures as evident in magnified unstained images **(Figure 3B)**, whole-mount skeletal stains **(Figure 3C**), and reconstructed microCT images **(Figure 3D)**. This recapitulates the craniosynostosis seen in humans with OGD.

**Figure 3.**
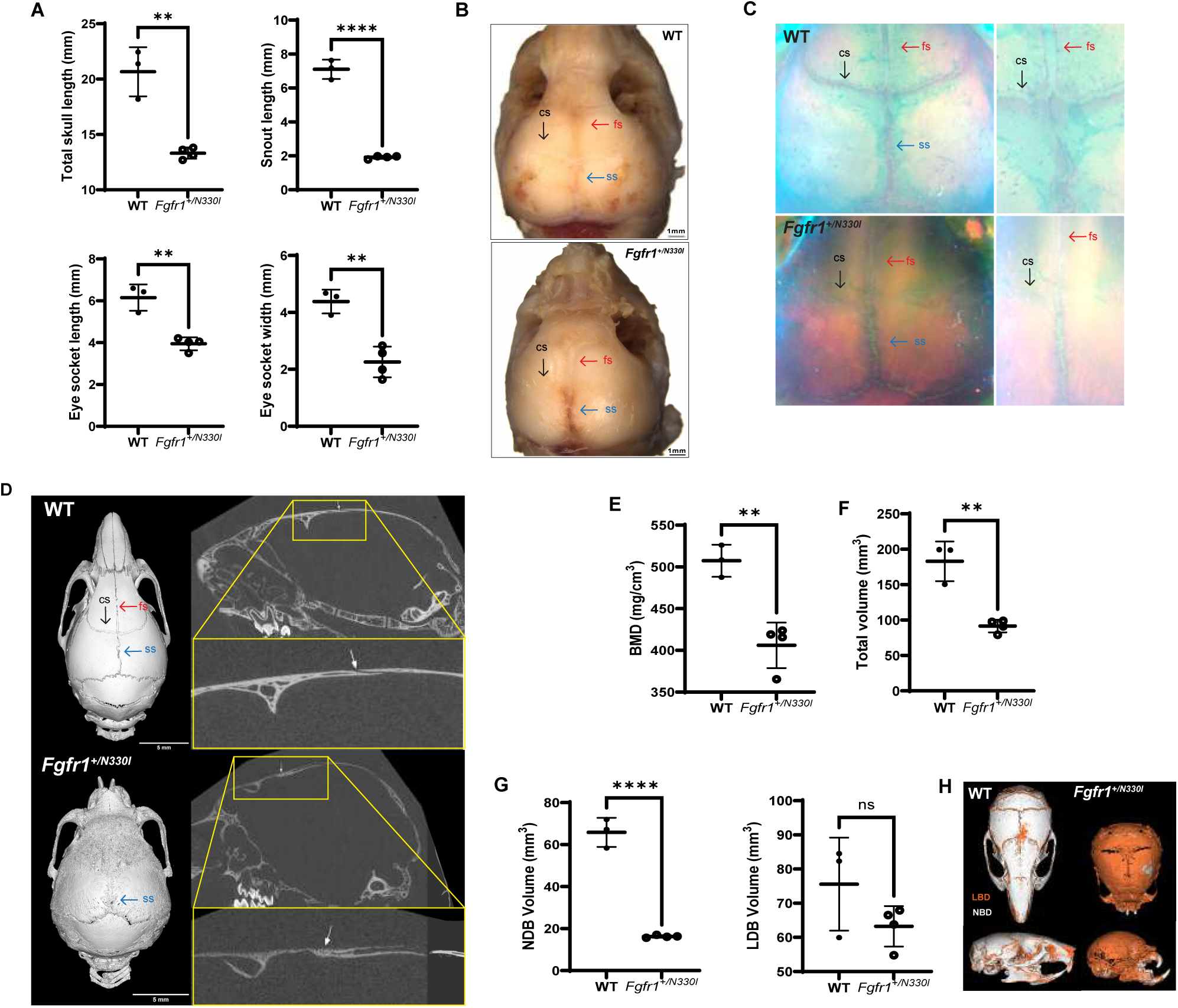
*Fgfr1^+/N330I^* mice show craniofacial involvement. **(A)** *Fgfr1^+/N330I^* mice show profound differences in skull morphology as shown by a reduction in total skull and snout length (35% and 72%) and reduction of the eye socket length and width (33% and 54%). Horizontal and vertical lines represent the mean ± SD of 3 WT or 4 mutant mice. **(B)** Representative images of unstained dissected calvaria from WT mice (top) and mutant (bottom) mice (scale bar = 1 mm) and **(C)** representative images from whole-mount skeletal stains show well-defined sutures in WT mice (top) whereas the mutant mice (bottom) show complete fusion of coronal and partial fusion of frontal sutures; magnified view of sutures in WT and mutant mice are shown on the right panel. fs-frontal suture, cs-coronal suture, ss-sagittal suture. **(D)** Representative 3D rendering from microCT scans of the skull (left), sagittal view (middle), and zoomed-in sagittal view (right), showing an open coronal suture in the wild-type (top panel) and a fused suture in the mutant (bottom panel). Resolution 10 µm, scale bar = 5 mm. **(E)** *Fgfr1^+/N330I^* mice have reduced skull BMD (20% reduction), **(F)** a smaller skull as shown by a marked reduction of its total volume, and **(G)** reduced volume of the bone with normal density (NBD, 78% reduction), but not of lower density bone (LDB), as compared to WT controls. Horizontal and vertical lines represent the mean ± SD of 3 WT or 4 mutant mice. **(H)** 3D reconstructions of WT and *Fgfr1^+/N330I^*skulls highlighting differences in bone density (NDB in white vs. LDB in orange). (ns = not significant, ** = p < 0.01, **** = p < 0.0001).

MicroCT revealed a 20% reduction of skull BMD in the mutant mice (**Figure 3E**). Additionally, the total skull volume was reduced by 50% (**Figure 3F**). Bone quality was analyzed by grouping different ranges of BMD; whereas the lower density bone (LDB, 200 - 599 mg/cm^3^) volume was comparable between the WT and mutant mice, the volume of normal density bone (NDB, ≥ 600 mg/cm^3^) in the mutant mice was 22% of that in WT mice (**Figure 3G**), highlighting the presence of a larger amount of undermineralized bone in the mutant mice (**Figure 3H**).

### Osteoglophonic dysplasia is a chondrodysplasia driven by impaired proliferation and apoptosis

Histology of 5-week-old mice hindlimbs revealed abnormal architecture of both the growth plates and secondary centers of ossification (SOC) (**Figure 4A and B**). Specifically, the growth plates showed abnormal seriation and asynchronous maturation of chondrocytes, no longitudinal septa, and accelerated terminal differentiation of hypertrophic chondrocytes. Further, *Fgfr1^+/N330I^* mice had delayed development or absence of secondary centers of ossification (**Figure 4B**). Moreover, we found differences in the distribution of proteoglycans both in the growth plates and articular cartilage (AC), further indicating disorganization of cartilage extracellular matrix (**Figure 4C**). These differences were also found in older mice (**Supplemental Figure S1**).

**Figure 4.**
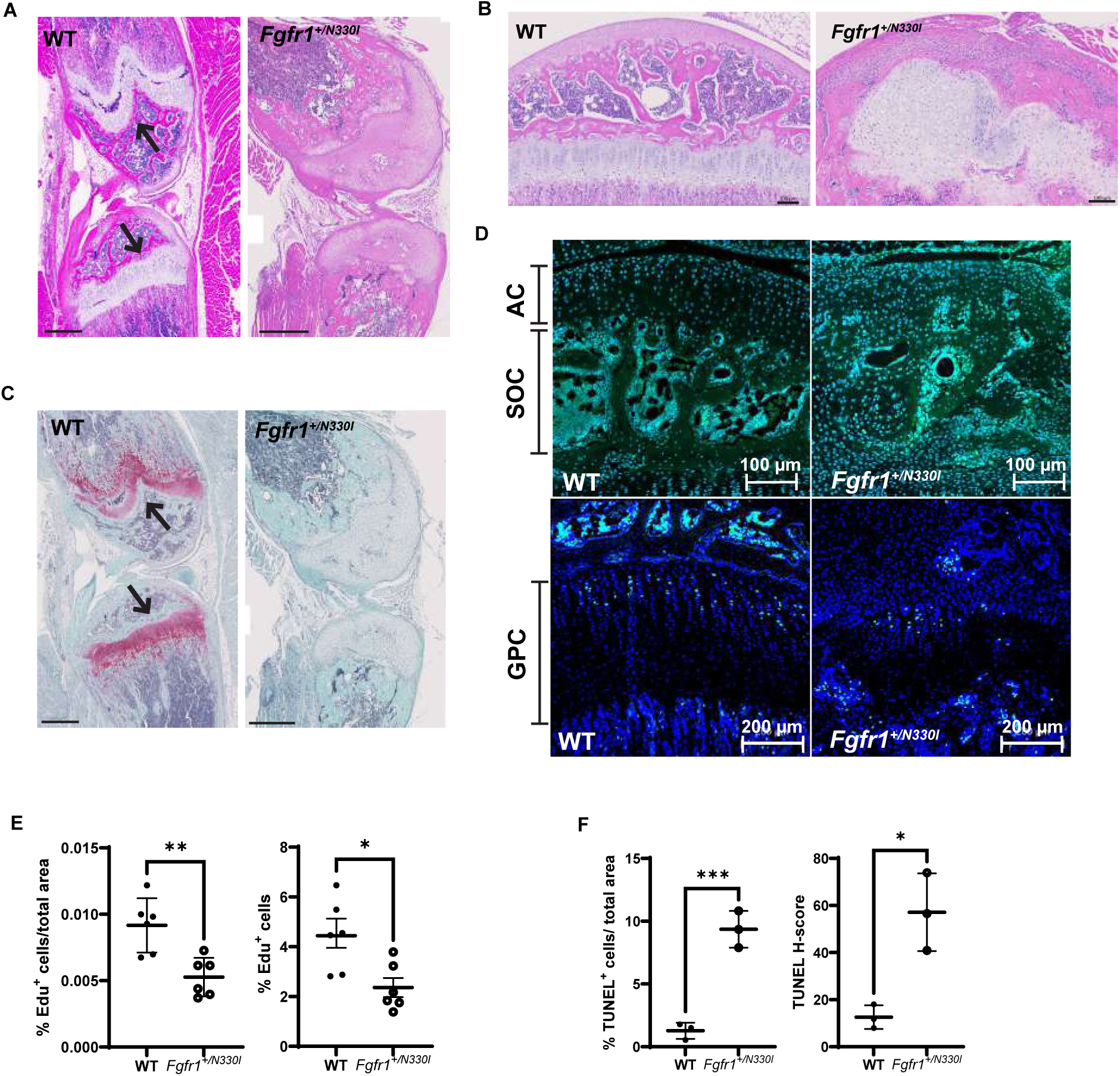
*Fgfr1^+/N330I^* mice display profound cartilage disorganization in femurs. **(A)** Qualitative histological analysis of Hematoxylin & Eosin (H&E) staining in 5-week-old *Fgfr1^+/N330I^* mice showing severe abnormalities of both growth plates (black arrows; scale bar = 500 µm) and **(B)** epiphyseal centers of ossification (scale bar = 100 µm). **(C)** Decreased Safranin O/Fast green (SafO/FG) staining for proteoglycans in mutant mice revealed altered composition of the cartilage extracellular matrix (red, black arrows). Scale bar = 500 µm **(D)** Representative images of cell proliferation assay in proximal tibial articular cartilage (AC) and growth plate cartilage (GPC) of mice representing nuclear staining with Hoechst 33342 (blue signal) and with EdU (green signal) revealed decreased proliferation in articular cartilage (AC) and secondary ossification center (SOC) in *Fgfr1^+/N330I^* mice when compared to WT mice (n = 2, 3-week-old). Scale bar = 100 µm (AC, SOC); scale bar = 200 µm (GPC). **(E)** Dot plots represent the percentage of EdU+ (left) and percentage of EdU+ cells normalized to tissue area in GPC (right), revealing significantly lower proliferation in *Fgfr1^+/N330I^* mice when compared to WT mice. Data points represent mean ± SD of three non-consecutive sections from two WT and two mutant mice. **(F)** Increased apoptosis demonstrated in femurs from mutant mice through quantification of TUNEL+ cells and relative H-score. Data points represent mean ± SD of 3 mice/group. (* p < 0.05, ** p < 0.01).

*Fgfr1^+/N330I^* mice showed a significant decrease (p = 0.01) in the percentage of proliferating chondrocytes in the growth plate of the proximal tibia when compared to their WT littermates (**Figures 4D and 4E**). This decreased rate of cellular proliferation was also observed in the epiphyseal region with severely disorganized AC and SOC. TUNEL staining was increased in femurs of *Fgfr1^+/N330I^* mice compared to those of WT controls (**Figure 4F**), indicating increased apoptosis.

Immunohistochemistry did not demonstrate significant differences in the levels of SOX9 (**Supplemental Figures S2A and S2B**) and Collagen X (COL10A1) (**Supplemental Figures S2C and S2D**), which represent markers of chondrocyte differentiation and hypertrophy, respectively. However, *Fgfr1^+/N330I^*mice showed reduced mRNA levels of osteoblast differentiation markers in cortical bone explants. Although we did not find significant differences in the levels of markers of osteogenic differentiation such as *Cbfa1* and *Col1a1* (**Supplemental Figure S3A**), the expression of *Spp1* and *Alpl*, encoding for osteopontin and alkaline phosphatase, respectively, were significantly downregulated in mutant mice (**Supplemental Figure S3B**). Consistent with the reduction of these critical mineralization markers, *Fgfr1^+/N330I^*mice also had reduced levels of *Tgfb* (**Supplemental Figure S3B**), an important factor for osteoblast differentiation (19).

### A pan-FGFR inhibitor, infigratinib, partially restores the size of Fgfr1^+/N330I^ mice

Since OGD is caused by an overactive FGFR1 receptor, we hypothesized that inhibition of the signaling pathway might lead to phenotypic improvement. We tested a small molecule pan-FGFR inhibitor (infigratinib) for ERK phosphorylation in cultured mutant bone marrow stromal cells (BMSCs) and for restoration of growth in our animal model of OGD. The half maximal inhibitory concentration (IC50), a quantitative measure of the potency of infigratinib at inhibiting MAPK phosphorylation in mutant cells, was calculated at 1.26 nM (**Figure 5A**), similar to the IC50 (0.9 nM) reported against purified recombinant FGFR1 under optimized conditions (20). Cultured BMSCs from *Fgfr1^+/N330I^*mice incubated with infigratinib for 1 hour exhibited a 76% decrease of kinase activity compared to vehicle control (**Figure 5B**).

**Figure 5.**
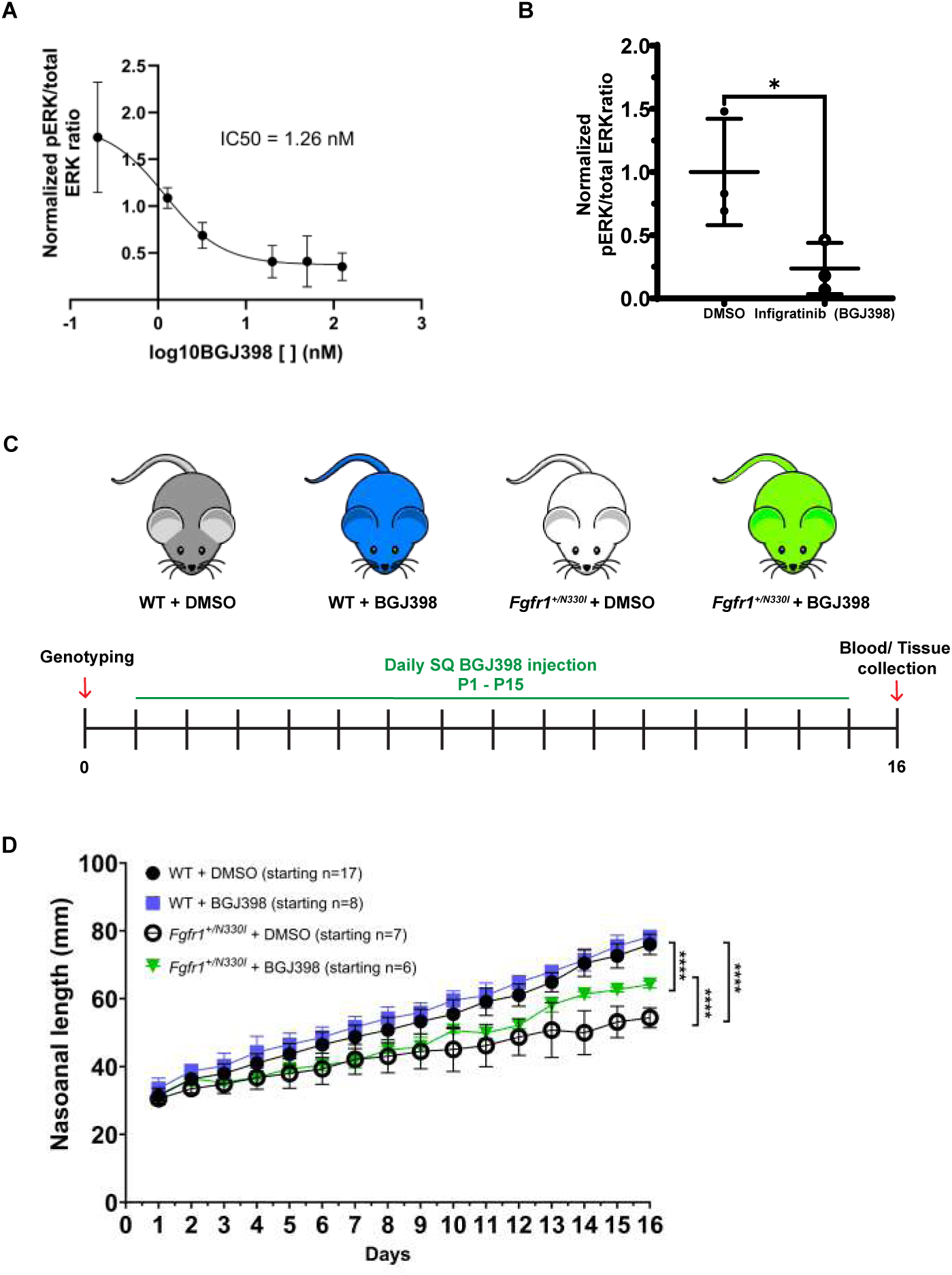
In vitro and in vivo evaluation of the effect of FGFR signaling pathway modulation on the phenotype of OGD. **(A)** Bone marrow stromal cells (BMSCs) from mutant mice were cultured in triplicate and a dose-response curve was obtained; the IC50, or the concentration of a substance (infigratinib) needed to inhibit a biological process (the phosphorylation of ERK) by 50%, was calculated at 1.26 nM. **(B)** BMSCs were isolated from *Fgfr1^N330I/+^* mice and cultured to ∼80% confluence; cells were then incubated with 125 ng/mL of infigratinib for 1 hour at 37°C. A 76% decrease in downstream phosphorylation compared to vehicle control was noted. Horizontal and vertical lines represent the mean ± SD of three biological replicates. * p < 0.05. **(C)** Graphical representation of the experimental design. The preclinical study involved genotyping on day of birth followed by subcutaneous injection of infigratinib 2 mg/kg/d of SQ infigratinib x 15 days to assess its effect in vivo. **(D)** Growth curve during infigratinib treatment trial. The naso-anal length of treated mutant animals (green inverted triangles) was intermediate between that of untreated mutant mice (empty circles) and WT mice (black circles and blue squares), with a significant difference in length between the treated mutant and the untreated (vehicle-treated) mutant mice (**** p < 0.0001 using a mixed model for repeated measures with time x treatment interaction effect). n = 6-17 mice/group.

To test the disease-modifying effect of infigratinib in vivo, all mice were genotyped at birth and treated with either vehicle (3% DMSO and 3.5 mM HCL) or drug (infigratinib). Daily subcutaneous injections were administered between postnatal days 1 and 15 (2 mg/kg/d of infigratinib or equivalent volume of vehicle) (**Figure 5C**). Infigratinib treatment of *Fgfr1^+/N330I^*mice showed no effect on mortality rate as indicated by the number of deaths by postnatal day 16 (3 of 7 vehicle-treated mice vs. 4 of 6 infigratinib-treated mice). However, infigratinib significantly improved the naso-anal length of *Fgfr1^+/N330I^* mice (**Figure 5D**).

### Fgfr1^+/N330I^ mice show impaired FGF23-phosphate axis metabolism and consequently impaired skeletal mineralization

In addition to the chondrodysplasia phenotype, we investigated the potential impact of FGFR1 p.N330I on phosphate wasting, which is a manifestation of OGD patients with this variant. Mutant mice (2-3 weeks old) exhibited 17-fold increased plasma levels of intact FGF23 compared to WT littermates (**Figure 6A**). In parallel with the FGF23 excess, *Fgfr1^+/N330I^* mice also exhibited significantly reduced phosphate when compared to their WT littermates (**Figure 6A**). These findings reflected severe dysregulation of FGF23 production and impairment of phosphate homeostasis and bone mineralization.

**Figure 6.**
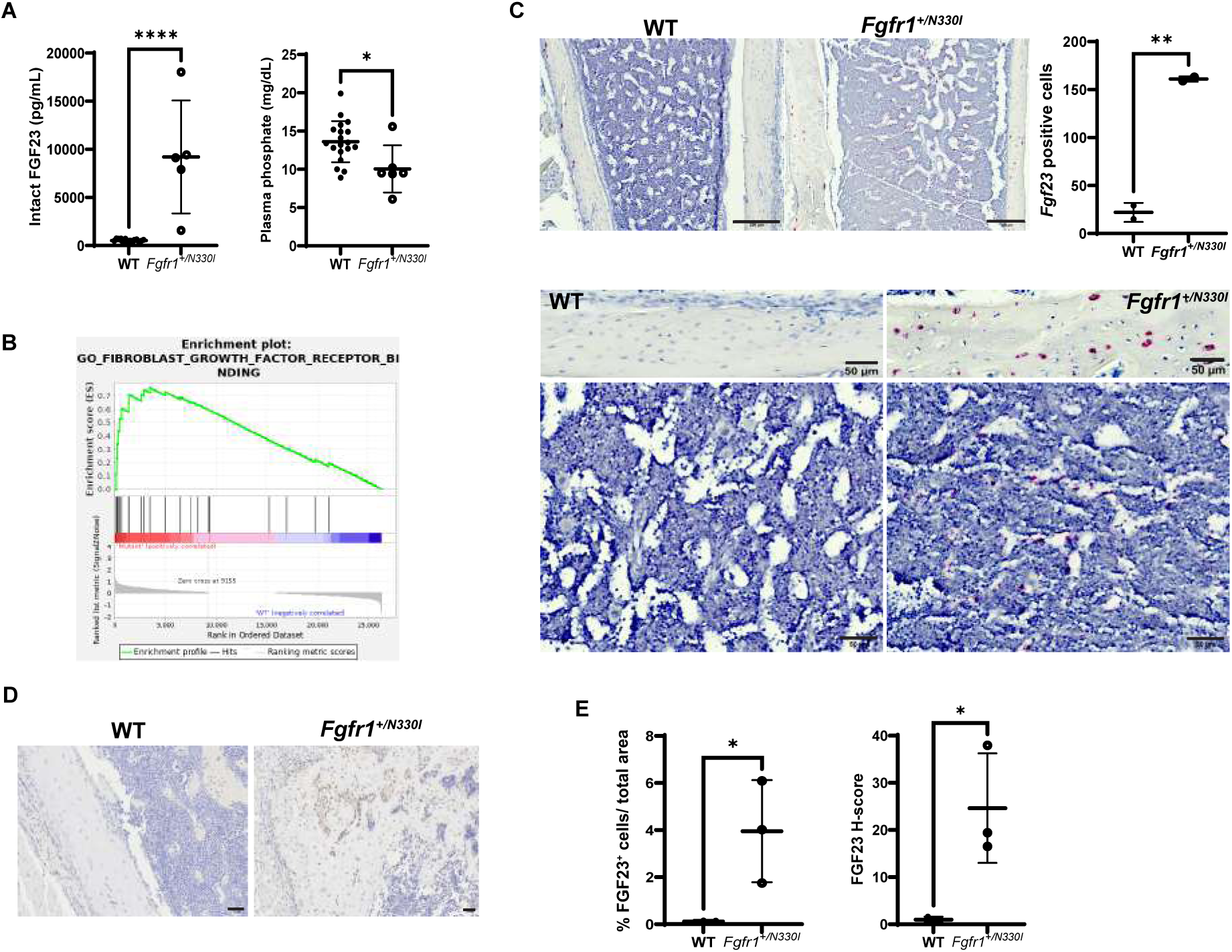
Effect of the Fgfr1 N330I variant on FGF23 synthesis. **(A)** *Fgfr1^+/N330I^* mice show markedly elevated intact FGF23 levels (9211 ± 2625 pg/mL; n = 5), when compared to WT littermates (522.1 ± 50.2 pg/mL; n = 14; p<0.0001) and consequently reduced circulating phosphate levels (10.1 vs. 13.6 mg/dL, p = 0.0115; n = 19 WT and 6 mutant mice). Horizontal and vertical lines represent the mean ± SD. **(B)** Transcriptomics in bone (n = 5) revealed enrichment of the FGFR binding pathway (FDR adjusted p-value = 0.036), with *Fgf23* having the highest rank metric score for the genes driving enrichment within this pathway. **(C)** Representative images of mouse femur sections processed for RNAscope in situ hybridization. *Fgf23* mRNA probe shows red staining while nuclear staining is blue. The top panels demonstrate femur at 20× magnification (scale bar = 200 µm), while the middle panels represent zoomed-in view of cortical bone (scale bar = 50 µm) demonstrating increased *Fgf23* staining in mutant femurs. Bottom panels represent zoomed in view of the marrow region (scale bar = 50 µm) showing *Fgf23* staining in peri-sinusoidal cells. Dot plot represents the density of positively stained nuclei with *Fgf23* probe with respect to the area of cortical bone. The mean number of cells staining for *Fgf23* in the mutant bone was 161, significantly higher in comparison to that of the WT mice (22, p = 0.0027). Horizontal and vertical lines indicate the mean ± SD from 2 WT and 2 mutant mice. **(D)** FGF23 protein levels were also increased in femurs of mutant mice as shown by immunohistochemical staining. Original magnification = 40×; scale bar = 100 µm. **(E)** Quantification of FGF23 abundance in the femurs of WT and mutant mice, highlighting not only a higher percentage of FGF23-positive cells but also stronger intensity indicating increased per-cell production (H-score) in the mutant mice as compared to WT controls (3 mice/group). Horizontal and vertical lines represent the mean ± SD of 3 mice/group.

FGF23 is predominantly produced by mature osteoblasts and osteocytes. We performed whole femur RNA-Seq transcriptomics to assess which pathways are dysregulated in the long bones of OGD mice. Gene Set Enrichment Analysis revealed enrichment of the FGFR binding pathway in mutant bone, with a Normalized Enrichment Score of 1.94. Genes driving FGFR enrichment included *Fgf23* (displaying the highest rank metric score of 0.962), *Fgf1*, *Fgf10*, *Fgf18*, *Fgf16*, *Fgf9*, *Fgf2*, and *Fgf22* (**Figure 6B**). In line with increased systemic levels, local FGF23 production was also significantly higher in the femurs of *Fgfr1^+/N330I^* mice. RNAscope in situ hybridization revealed prominent *Fgf23* staining in both the cortical and marrow region of femurs from *Fgfr1^+/N330I^*mice, further suggesting upregulation of *Fgf23* expression (**Figure 6C**). Whereas *Fgf23* expression in WT mice was predominantly restricted to the periphery of cortical bone, in mutant mice its expression was distributed throughout the bone cortex. *Fgf23* was also detected within the marrow space in a peri-sinusoidal fashion, suggesting that increased FGFR1 signaling induces *Fgf23* expression in skeletal stem cell pericytes present in the bone marrow. In cortical bone, 7 times more *Fgf23*-positive osteocytes were detected. Similarly, protein levels of FGF23 were higher in osteocytes and were also detected in osteoblasts and lining cells (**Figure 6D**). Specifically, the percentage of FGF23-stained cells normalized to the total area was 0.1% in WT mice and 4% in mutant mice. The histochemical scoring (H-score) assessment was also significantly higher (25% vs. 1% in mutant and WT mice, respectively) indicating that a higher amount of FGF23 per cell was produced (**Figure 6E**).

Given the increase in plasma FGF23 and decrease in plasma phosphate levels, we assessed whether these abnormalities in mineral metabolism translated into decreased mineralization of bone tissue. Histology of undecalcified bone samples confirmed the profound skeletal tissue disorganization and revealed also an increase in the amount of unmineralized bone matrix (osteoid) (**Figure 7A**). Indeed, by histomorphometry, the osteoid surface in femurs from *Fgfr1^+/N330I^* mice trended 2.5-fold higher than in WT mice, and the osteoid volume and width were significantly higher (4.3-fold and 1.8-fold, respectively) (**Figure 7B**). In line with this, the amount of mineralized bone as determined by von Kossa staining was significantly lower (**Figure 7C** and **7D**). Altogether, these findings corroborated the defective mineralization in the bone of mutant mice.

**Figure 7.**
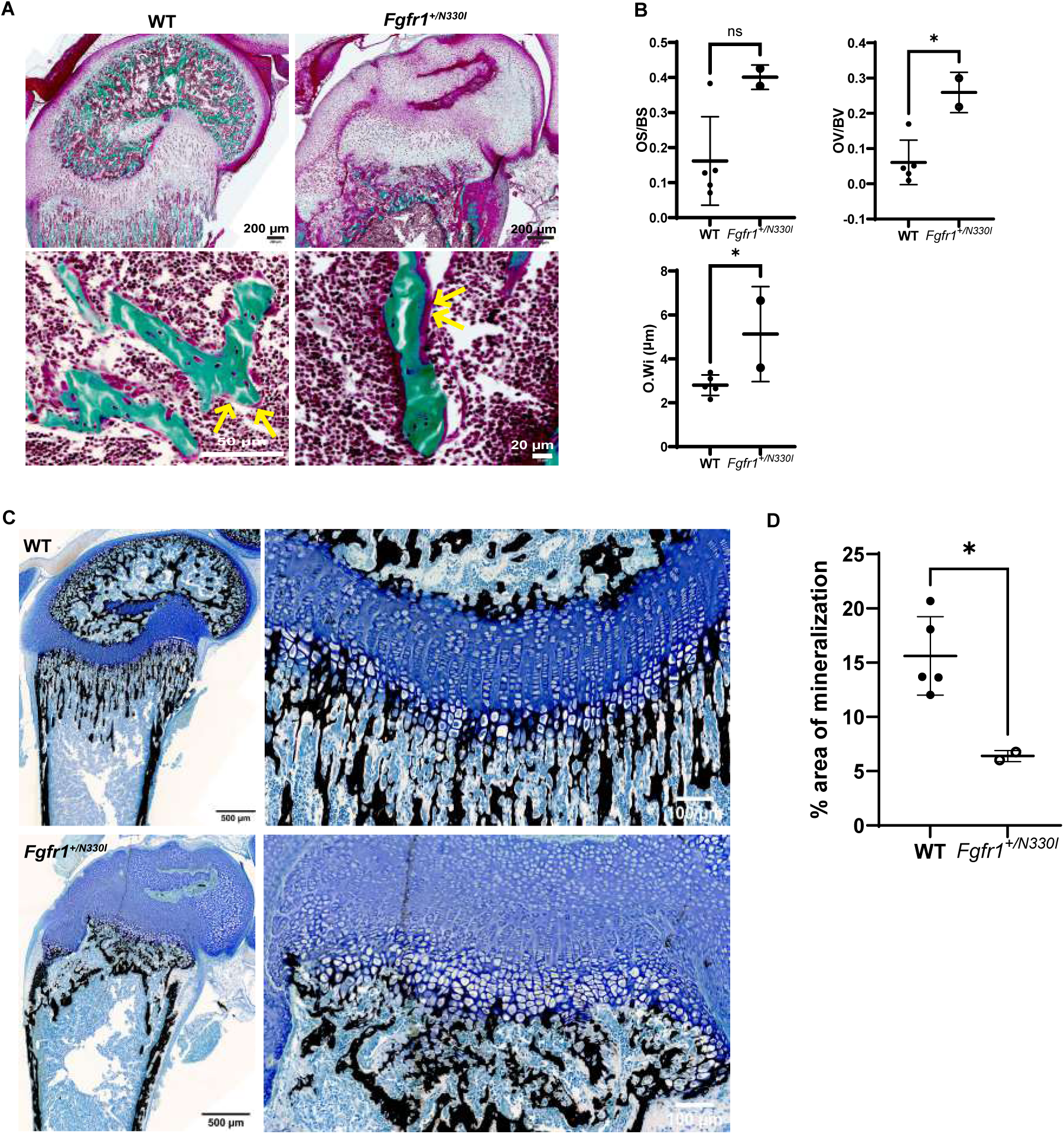
*Fgfr1^+/N330I^* mice show impaired mineral metabolism. **(A)** Representative images of undecalcified mouse femur sections stained with Goldner trichrome stain showing the impaired development and organization of the articular cartilage, secondary ossification center and growth plate (top panel) and revealing the increased osteoid (mineralized bone in green, and osteoid seam in red marked with yellow arrows; bottom panel) in *Fgfr1^+/N330I^* mice. **(B)** Dot plots representing static histomorphometry parameters that illustrate a trend towards increased osteoid surface fraction (40% vs. 16%, p = 0.054), and a significant increase in osteoid volume fraction (26% vs. 6%, p = 0.012) and osteoid width (5.1 mm vs. 2.8 mm, p = 0.046) in femurs from *Fgfr1^+/N330I^*mice. **(C)** Representative images of von Kossa-stained sections from the distal end of undecalcified femurs from WT and mutant mice showing mineralized tissue as black (left panels, scale bar = 500 µm). Brightfield images from anatomically equivalent regions of distal femurs were used to quantify the area of mineralization using ImageJ. Zoomed-in view of distal growth plate (right panels, scale bar = 100 µm). (D) Dot plot representing the percentage (%) of mineralized area. Data are presented as mean ± SD; p = 0.019. Horizontal and vertical lines represent the mean ± SD of 5 WT and 2 mutant mice. Original magnification 20×. OS-osteoid surface, BS-bone surface, OV-osteoid volume, BV-bone volume, O.Wi-osteoid width. (ns p ≥ 0.05, * p < 0.05, ** p < 0.001, **** p < 0.0001).

### Bone marrow organoids using Fgfr1^+/N330I^ osteoprogenitors show osteodysplasia and allow longitudinal evaluation of the FGF23-phosphate axis

The severe phenotype in the *Fgfr1^+/N330I^* mouse model typically led to lethality within the first two weeks of life. In addition, the small size of the animals limited blood sampling, hindering our ability to obtain longitudinal evaluations of mineral metabolism. To address these shortcomings, we used a well-established in vivo allograft assay, transplanting cultured BMSCs along with an appropriate scaffold into subcutaneous pockets in immunocompromised mice (in vivo ectopic ossicles) (21) (**Figure 8A**). We transplanted two million WT or *Fgfr1^+/N330I^*BMSCs depleted of hematopoietic cells into subcutaneous pockets of immunocompromised mice for 80 days, at which time the transplants were collected for microarchitectural evaluation. MicroCT analysis revealed that *Fgfr1^+/N330I^*BMSC transplants lack cortex, are larger (2.4-fold increase) and contain more bone (6.7-fold increase) than WT transplants (**Figure 8B, Supplemental Table ST2)**. Trabeculae are 3.5-fold more numerous, 1.6-fold thicker, 3.3-fold closer together and 5-fold more interconnected. However, *Fgfr1^+/N330I^* BMSC transplants have less bone surface (2.2-fold), the bone has less mineral density (11% decrease), and transplants have a dysmorphic shape, reflected by a structural model index (SMI) of -1.8, representing a high preponderance of concave structures (spheroids) over rods or plates. In contrast, the SMI in WT BMSC transplants was 2.3, reflecting a high abundance of trabecular plates. Lastly, trabeculae from *Fgfr1^+/N330I^* BMSC transplants were less organized spatially, as reflected by their lower degree of anisotropy (18% decrease) **(Supplemental Table ST2)**. Histology of the transplants via H&E staining was consistent with the microCT findings, revealing increased bone mass with lack of cortex (**Figure 8C**).

**Figure 8.**
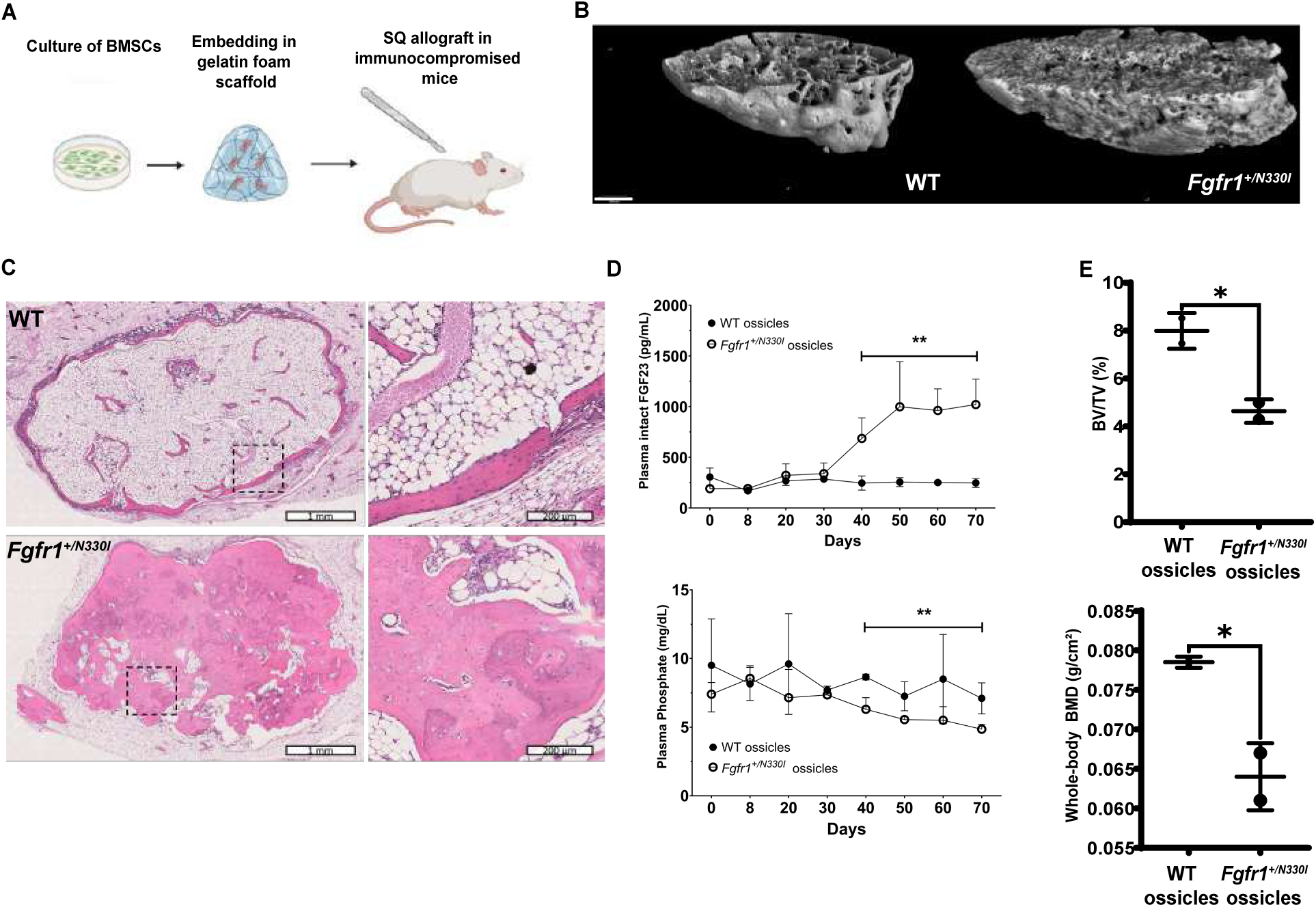
An in vivo ectopic ossicle model reveals osteodysplasia and allows longitudinal assessment of the disrupted FGF23-phosphate axis. **(A)** Schematic representation of the in vivo transplantation assay. Gelatin sponges were loaded with 2 million BMSCs (WT vs. *Fgfr1^+/N330I^*) depleted of hematopoietic cells and transplanted into subcutaneous pockets in immunocompromised mice for 80 days. Created with BioRender.com. **(B)** Representative 3D microCT reconstruction of in vivo transplants. Scale bar = 500 µm. **(C)** Low and high magnification H&E staining images of representative transplants loaded with WT BMSCs (top panel) and *Fgfr1^+/N330I^* BMSCs (bottom panels). The low magnification image for the *Fgfr1^+/N330I^* BMSC transplant was obtained by merging two consecutive section images from the scanned slide. Histology confirmed excess bone but lack of cortex in the *Fgfr1^+/N330I^* BMSC transplants. **(D)** Plasma intact FGF23 concentrations remained stable in mice receiving WT BMSC transplants, while those receiving *Fgfr1^+/N330I^* BMSC transplants showed elevation of intact FGF23 levels starting at 40 days post-transplantation (top). Similarly, plasma phosphate concentrations were decreased in mice receiving *Fgfr1^+/N330I^* BMSC transplants starting at 40 days post-transplantation (bottom). ** p < 0.01. **(E)** Decreased bone volume fraction as measured via microCT (BV/TV 4.64 vs. 7.99%; p = 0.03) and decreased whole-body BMD as measured via DXA (0.0640 vs. 0.785 g/cm^2^; * p = 0.04) in mice receiving *Fgfr1^+/N330I^* BMSC transplants compared to those receiving WT BMSC transplants. (* p < 0.05).

The in vivo ossicle model allowed us to evaluate systemic bone mineral metabolism in a longitudinal fashion. We obtained blood from the transplanted mice every 10 days by retro-orbital collection, and measured plasma levels of intact FGF23 and phosphate to assess systemic bone mineral metabolism. Plasma levels of intact FGF23 increased in mice receiving *Fgfr1^+/N330I^* BMSC allografts starting at 40 days post-transplantation (**Figure 8D**). Indeed, statistical comparisons of plasma intact FGF23 levels from days 0-30 revealed no differences (261 vs. 257 pg/mL; p = 0.93), whereas the same comparison from days 40-70 revealed significantly increased levels (917 vs. 250 pg/mL; p = 0.0032) in mice with *Fgfr1^+/N330I^* ossicles compared to mice with WT ossicles. Similarly, plasma phosphate levels remained unchanged between days 0-30 (8.7 vs. 7.6 mg/dL; p = 0.20), whereas between days 40-70 mice with *Fgfr1^+/N330I^* ossicles exhibited lower phosphate concentrations compared to mice with WT ossicles (5.6 vs. 7.9 mg/dL; p = 0.0032). Consistent with these findings, DXA analysis of mice at 80 days post-transplantation revealed a 19% decrease in whole body BMD and microCT analysis of femurs revealed a 42% decrease in bone volume fraction in mice receiving *Fgfr1^+/N330I^* BMSC transplants compared to those receiving WT BMSC transplants (**Figure 8E**). This strongly suggests that the expression of the *Fgfr1^+/N330I^* variant in bone drives FGF23 excess leading to hypophosphatemia, which in turn leads to systemic undermineralization of bone.

## Discussion

We created the first animal model of OGD, a disorder characterized by craniosynostosis, distinctive craniofacial features, profound short stature with femoral shortening, and osteopenia (9) caused by GoF variants in the Fibroblast Growth Factor Receptor 1 gene (*FGFR1*), resulting in constitutive receptor tyrosine kinase activity and aberrant downstream signaling. The *Fgfr1^+/N330I^* mouse harbors the *Fgfr1* c.989A>T (p.N330I) variant, which is associated with FGF23 excess in OGD. The mutant mice proved to be a true model of the human disease. They exhibited notably small size, as evidenced by decreased whole-body length, naso-anal length, and femoral length, along with distinct craniofacial features including craniosynostosis, recapitulating the human phenotype.

The murine model provided insights into the mechanisms of OGD. The long bones exhibited osteochondrodysplasia, with a disorganized growth plate, abnormal distribution of proteoglycans within the cartilage extracellular matrix, lack of secondary ossification centers within the epiphyses, and decreased cortical thickness with increased cortical porosity. We showed that this phenotype is driven at least in part by decreased proliferation of cells within the growth plate, articular cartilage and secondary ossification center, and by cell apoptosis.

The mutant mouse also allowed us to investigate the FGF23-phosphate axis as it related to excess FGFR1 signaling. Plasma phosphate was reduced in mutant mice compared to WT, associated with a marked elevation in circulating FGF23 levels. RNA-Seq analysis from long bones showed enrichment of the FGFR binding pathway in mutant cortical bone, with prominent overrepresentation of *Fgf23*. Further, in situ hybridization in mutant bones demonstrated abundant *Fgf23* expression throughout cortical bone and in bone marrow pericytes, contrasting with sparse expression confined to the subperiosteal cortex in WT mice.

A relevant topic is whether FGFR1 acts solely as an FGF23 receptor or also as an inducer of FGF23 expression and secretion. It seems counterintuitive that the activation of a receptor would lead to an increase in the amount of its ligand. FGF23 is a well-established ligand for FGFR1 (22); as for most other hormones, activation of FGFR1 by FGF23 would be expected to lead to decreased levels of ligand via a negative feedback loop. However, while FGF23 is synthesized and regulated in bone cells, FGF23 signaling occurs in the renal tubular epithelial cells, with downregulation of phosphate transporters leading to phosphaturia and inhibition of vitamin D activation leading to reduced calcitriol. Thus, there is a compartmental separation between the tissue of origin of FGF23 and its site of action with respect to mineral metabolism; the seemingly paradoxical increase in FGF23 ligand could be explained by the role of FGFR1 signaling in FGF23 synthesis. There is additional published in vitro, in vivo, and clinical evidence that FGFR1 signaling increases FGF23 production. In a mouse MC3T3-E1 osteoblastic cell line, FGFR1 signaling was shown to regulate *Fgf23* gene transcription via a MAPK dependent pathway and FGF23 protein expression via a PI3K-AKT depending pathway (23). Moreover, pharmacological activation of FGFR1 by monoclonal antibodies in adult mice led to increased serum FGF23 levels and mild hypophosphatemia, and the same monoclonal antibody in cultured rat calvarial osteoblasts induced Fgf23 mRNA expression and protein secretion into the culture medium (24). Clinically, phosphaturic mesenchymal tumors associated with FN1-FGFR1 fusion proteins cause increased FGFR1 signaling, resulting in ectopic FGF23 synthesis (25). Pharmacologic FGFR1 inhibition (such as with infigratinib) leads to reduced FGF23 and increased phosphate levels (26, 27). Our current work adds further evidence to the growing body of literature relating FGFR1 signaling with FGF23 synthesis.

Another question is whether the FGF23 excess is driven by simple receptor activation or by some alternative mechanism. To this point, any of five known FGFR1 activating variants lead to the osteochondrodysplastic phenotype of OGD, but only certain variants

(p.Asp330Ile) lead to FGF23-mediated hypophosphatemia (9, 28). Thus, there appears to be a dissociation between osteochondrodysplasia (driven by any variant leading to FGFR1 activation) and FGF23 excess (driven by only specific activating variants) in OGD. FGFR1 is not only expressed in the plasma membrane, but is also translocated to the nucleus, and this nuclear translocation is dependent on the transmembrane domain (29). It was recently shown that N-glycosylation (addition of sugar chains into asparagine residues) of FGFR1 acts as a switch for nuclear translocation, so that glycosylation-deficient mutant FGFR1 is predominantly localized in the nuclear envelope (30). It is possible that our mouse model and patients with the corresponding variant experience FGF23 excess due to increased nuclear translocation of mutant FGFR1. First, variants leading to OGD are localized in or adjacent to the transmembrane domain, a region instrumental in nuclear translocation of the receptor (29). Second, p.Asp330, the variant amino acid residue in our *Fgfr1^+/N330I^* mouse model of OGD, is a known site of FGFR1 N-glycosylation, and deficient glycosylation due to substitution of the Asp residue should lead to increased nuclear localization of the receptor (30). Third, nuclear translocation of FGFR1 has been shown to upregulate FGF23 gene expression (31). Thus, it is possible that some variants leading to FGFR1 activation could also differentially lead to FGF23 excess.

Since the osteochondrodysplasia phenotype is driven by overactivation of the FGFR1 receptor, we evaluated the effect of infigratinib, a small molecule tyrosine kinase inhibitor against FGFR1-3 activity in vitro and in vivo. In cultured mutant BMSCs, infigratinib led to a marked decrease in kinase activity after 1-hour incubation compared to vehicle control, whereas mutant mice treated with infigratinib from postnatal day 1 to 15 showed improvement in naso-anal length when compared to untreated mutant mice. Results from this preclinical study have identified infigratinib as a promising option for OGD treatment. Infigratinib was used safely in a phase 2 study in children 3-11 years old with achondroplasia, a disorder caused by overactivity of the FGFR3 receptor (32), and is currently in phase 3 trials in children 3-18 years old with the same disorder (ClinicalTrials.gov ID: NCT06164951). Infigratinib could potentially be repurposed as a targeted therapy for patients with OGD, given the marked morbidity of this disease, i.e., profound short stature with skeletal deformities, osteopenia, bone pain, impaired mobility and recurrent fractures (28).

Constitutive mutant mice were not available because of infertility and decreased survival. *Fgfr1^+/N330I^* mice are virtually infertile; mutant females cannot carry a pregnancy to term, and mutant males cannot mount WT females. We successfully knocked the *Fgfr1^N330I^* variant into a mosaic founder that transmitted the allele to approximately 15% of its progeny. However, sperm harvesting from 12-week-old germline mutant males was unsuccessful due to the diminutive size of the animal and testicles. Therefore, experimental animals we generated were obtained via in vitro fertilization using sperm from the original mosaic founder.

Despite the shortcoming due to the animal infertility, the in vivo ossicle model enabled us to evaluate systemic bone mineral metabolism in a longitudinal fashion. Even though the transplants were decalcified and, therefore, did not allow us to determine the mineralization of the newly formed bone within the ossicles, our datasets clearly indicated that the *Fgfr1^+/N330I^* variant leads to skeletal dysplasia and impaired bone metabolism.

In conclusion, we developed the first animal model of OGD that closely recapitulates the skeletal phenotype observed in humans. Mineral metabolism studies in our model demonstrate excess FGF23 resembling that seen in patients with the same genetic variant. Our preclinical studies indicate that infigratinib is a promising drug for the treatment of OGD. We plan to establish a conditional inducible *Fgfr1* model that enables activation of mutant gene expression through Cre-mediated recombination. This approach will allow us to increase the number of animals available to study the underlying pathomechanisms of the disrupted FGFR1-FGF23-phosphate axis, contributing further insights into OGD as well as other bone and mineral disorders associated with FGF23 dysregulation. It will also facilitate the evaluation of infigratinib and related candidate drugs to address an unmet medical need for patients with OGD.

## Methods

### Sex as a biological variable

Both male and female mice were used for the experiments described here. Female and male mice between 2-5 weeks of age were used in all experiments. The animals were housed in filter-top micro-isolator standard cages on positive-pressure ventilated racks. Mice were provided ad libitum chlorinated and UV-treated RO water and an autoclaved or irradiated mouse diet.

### Generation of Fgfr1^+/N330I^ mice

*Fgfr1^em1Cferr^* (abbreviated *Fgfr1^+/N330I^*) knock-in mice modeling the human *FGFR1* c.989A>T (p.N330I; NM_023110.3; rs121909632) variant were generated using CRISR-Cas9-mediated homology-directed repair (HDR). A mix of guide RNA (GGAAATGGAGGTGCTTCATCTA, synthesized as IDT Ultramer RNA Oligo), a donor DNA repair template (1 µM each), and saCas9-KKH variant (33) was used for electroporation into FVB/N × C57BL/6J F1 hybrid eggs. The repair template (full sequence in Supplemental Data S1) was synthesized as IDT Megamer single-stranded DNA fragments with 250 bp and 248 bp isogenic homology arms flanking a portion of the 501-bp template reverse complement sequence is provided here: 5′…CCCGCATCCTCAAAGGAGACAaTgCGTAGATGAAGCACCTCCATT..3′ (lower case indicates the two nucleotide changes compared to wild-type). The resulting allele, *Fgfr1^em1Cferr^*, also referred to as *Fgfr1^+/N330I^*, contains two nucleotides different from wild-type (GRCm38 chr8:25,564,388-25,564,393 CGGAAT > CGcAtT) expected to result in a single amino acid change in exon 8 of NM_010206 (mouse FGFR1 p.Asn330Ile; equivalent to human FGFR1 p.Asn330Ile).

Sanger sequencing was used to confirm the knocked-in allele in founder animals (forward primer: 5’-CTCTCGGAGGTCTCATGTCC-3’; reverse primer: 5’-GAGCAGAAAAGGGTACGCAG-3’). Indels in the founder animals were ruled out by fluorescent PCR (forward primer with M13F tail: 5’-TGTAAAACGACGGCCAGTCTCTCGGAGGTCTCATGTCC-3’; reverse primer with PIG tail: 5’-GTGTCTTGAGCAGAAAAGGGTACGCAG-3’; universal M13F-FAM primer: 5’-FAM-TGTAAAACGACGGCCAGT-3’) (34). For genotyping experimental animals, DNA was extracted from pup tail biopsies and DNA prepared using a rapid DNA extraction kit (REDExtract-N-Amp Tissue PCR Kit; Sigma-Aldrich, MO, USA). Samples were amplified using a TaqMan Fast Advanced Master Mix for qPCR (ThermoFisher Scientific, MA, USA) with a VIC-labeled probe for wild-type sequence and a FAM-labeled probe for alternate sequence (ThermoFisher Assay ID: ANDKGJG). An ABI StepOne Plus instrument was used for thermocycling and detection.

### Morphometric measurements

Wild-type (WT) and mutant mice were anesthetized under isoflurane (1-3% flow rate) prior to recording various morphometric parameters. Weights were measured using a Sartorius BP3100S balance (readability: 0.01 g), and lengths were measured using Reed R7400 digital caliper (accuracy: 0.03 mm).

### Craniosynostosis analysis

The whole dissected calvaria from 3-week-old mutant (*Fgfr1^+/N330I^*) and wild-type (*Fgfr1^+/+^*) mice were rinsed with 1X PBS and stored in 70% ethanol to maintain tissue integrity. The calvaria were first scanned with a Scanco µCT 50 (SCANCO Medical AG, Bruttisellen, Switzerland) for assessment of sutures and images were reconstructed with Analyze 14.0 software (AnalyzeDirect, Inc., USA). For differential staining of cartilage and bone, we performed whole-mount calvarium staining with Alcian blue and alizarin red, using the method previously described by Rigueur and Lyons (35). Briefly, the calvaria were placed in acetone overnight followed by submerging in 0.03% Alcian blue stain (in 80% ethanol and 20% glacial acetic acid) overnight at room temperature (RT). The samples were destained by washing in 70% ethanol and leaving in 95% ethanol overnight. The solution was then replaced with 1% potassium hydroxide (KOH) for 1 hour followed by incubation in 0.005% Alizarin red stain (in 1% KOH) for 3 days at RT. The samples were again cleared with 1% KOH and placed in 100% glycerol until imaging. Following staining, cartilage stains blue by Alcian blue, whereas the mineralized bone stains red by Alizarin red. The images were captured using a digital camera mounted on a Zeiss SteREO Discovery.V12 microscope (Carl Zeiss AG, Oberkochen, Germany).

### In vivo transplantation

Two male *Fgfr1^+/N330I^* and two male WT littermates were euthanized at 4 weeks of age, and their femora and tibia were dissected under sterile conditions to establish BMSC cultures following previously established methods (36). Negative immunoselection using CD45-coated magnetic beads was performed 1-3 days before transplantation. Three transplants of two million WT or *Fgfr1^+/N330^*BMSCs per culture were loaded on a gelatin foam scaffold (Surgifoam, Ethicon, NJ, USA) and implanted in bilateral subcutaneous dorsal pockets in NSG immunocompromised mice (The Jackson Laboratory stock #005557, ME, USA) following previously established methods (37). Every 10 days, 0.1 mL of blood was obtained from the transplanted mice by retro-orbital collection, and plasma was processed for intact FGF23 and phosphate determinations. At day 80, mice were euthanized and fixed by perfusion.

### MicroCT analysis of mouse bones

Femur and skull bones from *Fgfr1^+/N330I^* mice and littermate controls were dissected and fixed with Z-fix fixative (Anatech, MI, USA) for 24 h, then rinsed twice with 1X PBS, pH 7.4 (Thermo Fisher Scientific) and stored in 70% ethanol at 4°C. Scans were obtained with a SCANCO µCT 50 machine and ex vivo microCT analyses were performed with the BMA add-on module of the Analyze 14.0 Software (AnalyzeDirect, KS, USA). The region of interest (ROI) was set using a manually-determined global threshold. Local BMD and three-dimensional microstructural cortical bone properties, including the bone volume fraction (BV/TV), cortical thickness (Ct.Th), cortical volume, cortical bone area fraction (Ct.Ar/Tt.Ar), cortical porosity (Ct.Po) and pore numbers (Po.N) were calculated according to the manufacturer’s software. After determining local BMD different density regions were identified: low density bone (LDB) (200 - 599 mg/cm^3^) and normal density bone (NDB) (≥ 600 mg/cm^3^)

For the NSG immunocompromised mice receiving the BMSC transplants, dissected femurs were preserved in gauze soaked with PBS-CaCl_2_ solution (PBS with 0.5 mM CaCl_2_) and stored in cryovials at -20°C until analysis. MicroCT images were acquired from the right femurs using a SkyScan 1174 scanner (Bruker Scientific, MA, USA) at 50 kV, 800 μA, and with a 0.25 mm Al filter, at a resolution of 6.8 μm/pixel. Scanning was performed with a 360° rotation, 0.6° angle increments, and two-frame averaging. A quartz piece was placed in the sample holder and calibrated to 0.801 g/cm³ mineral density by scanning alongside mouse bone phantoms. Each femoral scan was recalibrated using the quartz standard and surrounding PBS-CaCl_2_ solution, which matched the intensity of soft tissue (0 g/cm³). Images were reconstructed at 6.8 μm isotropic voxel size with 30% beam hardening correction using NRecon Software (Bruker) and analyzed with CTAn software (Bruker). Trabecular regions of interest (ROIs) were located adjacent to the distal femoral growth plate, extending proximally by 10% of the femoral length. The trabecular bone threshold was set at 0.250 g/cm³, based on mineral density profiles across multiple samples. Cortical ROIs were 10% of femoral length, centered between the third trochanter and distal growth plate. The cortical bone threshold was set at 0.5 g/cm³, similarly optimized. Analysis of selected left femurs revealed no significant differences compared to contralateral right femurs beyond the microCT measurement error.

### MicroCT imaging of BMSC transplants

BMSC transplants were scanned by microcomputed tomography (µCT 50, 70 kVp, 85 µA, 0.3 s integration time, 7 µm^3^ voxel resolution, Scanco Medical AG, Brüttisellen, Switzerland) and analyzed using the Scanco application center 1.0.28.0 software. For 3D reconstructions, a threshold of 443 mg HA/cm^3^ was used. Trabecular parameters and tissue mineral density (TMD) were calculated.

### Dual-energy X-ray absorptiometry (DXA)

An UltraFocus DXA cabinet (Faxitron®, AZ, USA) was used to measure whole body bone mineral density (BMD) and bone mineral content (BMC) of WT and *Fgfr1^+/N330I^* mutant mice. Following the calibration procedure as per the manufacturer’s manual, X-ray scans were imaged at 4× magnification.

DXA analysis of mice transplanted with WT or *Fgfr1^+/N330I^*BMSCs was performed using a Kubtec XPERT 80 system (Kubtec Medical, CT, USA). Total body DXA scans were performed at automatically selected exposures that ranged between 41.0 and 43.0 Kv, and total body (excluding head), femurs and L5 vertebrae bone mineral density and content (BMD, BMC) were analyzed using Kubtec Digimus software.

### Histology and immunohistochemical staining

Tissue samples from WT and *Fgfr1^+/N330I^* mice were fixed in Z-fix solution (Anatech) for 24h, decalcified in EDTA and subsequently embedded in paraffin. Longitudinal sections (5 µm thick) of long bones and vertebrae were stained with hematoxylin and eosin (H&E) and Safranin O/Fast Green (SafO/FG) for qualitative assessment of bone and cartilage architecture. Long bones were stained for FGF23 (1:1000, R&D, catalog #MAB26291), SOX9 (1:250, MilliporeSigma, catalog #AB5535), COL10A1(1:1000, Abcam, catalog #ab58632), and TUNEL. Slides were subjected to antigen retrieval in a sodium/citrate buffer (pH 6) for COL10A1 and FGF23, and Proteinase K solution for SOX9. Subsequently, slides blocked against endogenous peroxidases, and incubated with the primary antibodies. The antibodies were detected with a horseradish peroxidase (HRP)-conjugated system appropriately matched to the host of each primary, and visualized with DAB. Finally, the slides were counterstained with hematoxylin, dehydrated in graded alcohols, cleared in xylene, and mounted with permanent mounting media. All rinses were performed using either distilled water or TBST. For the TUNEL assay, slides were pretreated with Proteinase K, followed by EDTA, deionized water wash, and BSA blocking. Next, slides were incubated in a reaction mixture of TdT, dUTP, and buffer. Slides were then washed with SSC buffer and incubated with anti-digoxigenin antibody. Slides were visualized with ImmPRESS Vector Red (Vector Laboratories, CA, USA) and counterstained with hematoxylin. The slides were dehydrated with graded alcohols, cleared in xylene, and mounted in Permount mounting medium. Tissue embedding, all sectioning and stainings were performed by Histoserv Inc. (Germantown, MD, USA) using standard protocols for H&E and SafO/FG. Immunohistochemical stainings were quantified using QuPath v 0.5.1 (38). H-scores were calculated based on three intensity thresholds to categorize cells as *Negative*, *1+, 2+* and *3+* (weak, moderate or strongly positive staining, respectively).

### Static histomorphometry of undecalcified samples

Mouse femur and tibia with intact knee were dissected and fixed in 4% paraformaldehyde. Undecalcified bone samples were then processed as per routine procedure and embedded in methyl methacrylate, frozen and sectioned. Sections were stained with Goldner Trichrome and von Kossa as per standard procedures. Images were acquired using a Zeiss Axioscan slide scanner (Carl Zeiss AG) at 20× magnification for von Kossa-stained samples and at 40× magnification for trichrome-stained sections. Image analysis was performed using OSTEO v18.2.6 (Bioquant Image Analysis Corp., TN, USA). Von Kossa-stained sections (6 WT and 2 *Fgfr1^+/N330I^*mice) were used for structural indices, and trichrome-stained sections (5 WT and 2 *Fgfr1^+/N330I^* mice) were used for osteoid indices.

### 5-ethynyl-2’-deoxyuridine (EdU) fluorescent staining quantification

In vivo labeling of skeletal tissues with EdU was performed using the Click-iT® EdU Alexa Fluor® 488 Imaging Kit (Invitrogen catalog #C10337) as per manufacturers’ instructions. A 10mM stock solution of EdU (component A) was diluted in 2 mL of DMSO (component C), and 10 μL of EdU solution per g of mouse body weight was administered intraperitoneally into WT and *Fgfr1^+/N330I^* mice (3-week-old, n = 2/cohort); labelling was allowed for 2 hours and after euthanasia, femurs and tibias were collected. The tissues were fixed in 4% paraformaldehyde, decalcified in 14% EDTA in PBS (for 2-3 weeks at 4°C), embedded in paraffin, and 6µm longitudinal sections were mounted on slides; three non-adjacent sections from each mouse were used for EdU detection. Counterstaining was done with Hoechst 33342 nuclear stain and mounting with Vectashield medium. To assess the proliferation index in the growth plate, adjacent images comprising the entire growth plate of the proximal tibial region were captured using a Zeiss LSM 880 with Airyscan confocal microscope (Carl Zeiss AG) followed by quantitative analysis using Image-Pro v11 (Media Cybernetics, MD, USA). Both the total number of nuclei and EdU+ nuclei were counted. Whole tissue images were also digitized using a Zeiss Axioscan slide scanner with 20× objective (Carl Zeiss AG). A region of interest spanning the epiphyses and the entire growth plate was selected and divided into equal segments, with each segment analyzed for the average percentage of EdU+ nuclei.

### RNA isolation and RT-qPCR

Cortical bone explants were obtained from mouse femurs and total RNA was isolated with TRIzol reagents (Thermo Fisher Scientific). mRNA concentration was measured with a Nanodrop spectrophotometer and reverse-transcribed into complementary DNA (cDNA) using the High-capacity RNA-to-cDNA kit (Thermo Fisher Scientific). RT-qPCR was performed using the Applied Biosystems QuantStudio™ 7 Flex Real-Time PCR System (Thermo Fisher Scientific). Specific primer sequences used to assess gene expression are listed in **Supplemental Table ST1**. Glyceraldehyde 3-phosphate dehydrogenase (*Gapdh*) was used as housekeeping gene. Samples were normalized to the expression of *Gapdh* by calculating the comparative threshold: −ΔCt = − (C_t_ *gene of interest* − C_t_ *Gapdh*).

### RNA-Seq from bone

Femurs were collected from 3- or 4-week-old mice (5 *Fgfr1^+/N330I^*mice and 5 WT littermate controls). Total RNA from pulverized bone was extracted using TRIzol Reagent (Thermo Fisher Scientific) and further purified by phenol-chloroform precipitation. Total RNA (10 μg) was subjected to rRNA depletion–based library preparation and sequenced on an Illumina NovaSeq 6000 system (Active Motif, CA, USA). Between 26.5 and 32.3 million unique paired-end reads were generated for each sample. The alignment was performed using STAR v2.5.2b aligner (39) with mm10 as the reference genome. Data normalization was performed by the median of ratios method using DESeq2 v.1.14.1 (40), and normalized gene counts were used to perform Gene Set Enrichment Analysis (GSEA v4.0.3) (41) with default settings to determine whether members of a priori defined gene sets based on biological knowledge, i.e., genes sharing the same Gene Ontology (GO) category, are enriched.

### Plasma phosphate and FGF23 measurement

Retro-orbital blood collection was performed with heparinized micro-hematocrit capillary tubes (Fisherbrand catalog #22-362-566) and plasma was separated by centrifugation. The plasma samples were stored at -80°C until analysis. The inorganic phosphate concentrations for 18 WT and 6 *Fgfr1^+/N330I^*mice were measured either by IDEXX BioAnalytics or by the Division of Veterinary Resources at the NIH. Intact FGF23 levels were measured using a commercially available ELISA kit (Quidel, catalog #60-6800). Plasma aliquots of 25 µL were used for the assay as per manufacturer’s instructions.

### Fgf23 bone in situ hybridization

The tissue-specific mRNA expression of *Fgf23* was assessed with RNAscope 2.5 HD Reagent Kit – RED (ACD, catalog #322350) as per manufacturer’s instructions with slight modifications. Whole legs were collected from 3-week-old mice (n = 2 for each cohort) and were fixed in 4% paraformaldehyde in PBS for 3 days followed by decalcification in 14% EDTA at 4°C for 3-4 weeks. The decalcified samples were then embedded in paraffin and 5 µm longitudinal sections were used for the assay. Briefly, the slides were baked on a slide warmer at 60°C for 1h, deparaffinized in xylene, rehydrated in a graded series of ethanol solutions, and air-dried for 5 min. The samples were pre-treated with hydrogen peroxide for 10 min at room temperature and then with Custom Pretreatment Reagent (ACD, catalog #300040) for 15 min. After washing twice with deionized water, samples were completely air-dried, and hydrophobic barriers were drawn using PAP pen. Target probe (Mm-Fgf23), positive control probe (Mm-Ppib) and negative control probe (DapB) were added on the sections and hybridization was carried out in an oven at 40°C for 2 hours. After washing the probes, amplification was carried out using a series of Amp solutions, Amp1-Amp6, as per kit’s manual. After washings, slides were incubated with red solution at room temperature for 10 min, counterstained with 50% hematoxylin and mounted with EcoMount mounting media (Biocare Medical, CA, USA). After overnight air-drying, slides were digitized using a Zeiss Axioscan slide scanner (Carl Zeiss AG) at 20× magnification. Multiple ROIs from cortical bone comprising nearly all the cortical bone were considered and positive nuclei with red stain (>10 spots/nuclei) were quantified. Data are expressed as number of positively stained cells normalized to ROI area.

### Treatment with infigratinib phosphate

For the in vitro assessment of ERK phosphorylation, bone marrow stromal cells from *Fgfr1^+/N330I^* mice were cultured in duplicate at 37°C and 5% CO_2_ in a humid environment. Cells were maintained in growth media (αMEM with 10% non-heat inactivated fetal bovine serum/FBS, 1% glutamine, and 1% penicillin/streptomycin). The day before the experiment, cells were cultured in FBS-free media for overnight serum starvation. On the day of the experiment, cells were incubated for 1 hour with infigratinib in DMSO (10mM stock solution diluted with αMEM growth media without FBS), or the same dilution of DMSO control. The concentrations of infigratinib used for calculation of the half-maximal inhibitory concentration (IC50) were 125 nM, 50 nM, 20 nM, 3.2 nM, 1.3 nM and 0.2 nM. Cells were lysed with FastScan™ ELISA Cell Extraction Buffer (Cell Signaling Technology catalog #69905) with a protease/phosphatase inhibitor cocktail (Cell Signaling Technology catalog #5872); total ERK (Cell Signaling Technology catalog #67404) and phospho-ERK (Cell Signaling Technology catalog #42173) were measured by ELISA, following the manufacturer’s instructions. The phospho-ERK values were normalized to total ERK, to obtain normalized pERK/total ERK ratios. The concentrations of infigratinib were transformed to log10 scale (X = log10X); these log10 concentrations of infigratinib and the normalized pERK/total ERK ratios were used to calculate the IC50, using the GraphPad Prism built-in equation log(inhibitor) vs. response - Variable slope (four parameters).

The design of the in vivo preclinical trial involved genotyping the mice on the day of birth and allocating mice to separate cages for treatment with vehicle (DMSO) or drug (infigratinib). Non-pharmaceutical grade infigratinib phosphate was obtained from MedChemExpress (catalog #HY-13311A) as a 10 mM solution in 1 mL DMSO, aliquoted in small volumes of 20µL and stored at -20° Celsius. On the day of injection, this 20 µL stock solution was thawed and diluted with 617.5 µL of sterile saline (Fresenius Kabi, catalog #918610) and 22.5 µL 0.1M HCL (Sigma-Aldrich, catalog #2104) for a final concentration of 0.2 µg/µL (303 µM) infigratinib phosphate, 3% DMSO and 3.5 mM HCL. The dilution was performed using aseptic technique and filtered through a 0.22-micron syringe filter (Argos Technologies, catalog #FE12S). Animals were injected at a dose of 2 mg/kg (10 µl/g body weight) subcutaneously daily for 15 days using a 31-gauge needle (Brandzig, catalog #CMD2582). Another cohort of animals was injected with the same volume of vehicle, prepared using 20 µL DMSO (Sigma-Aldrich, catalog #276855), 617.5 µL of sterile saline, and 22.5 µL 0.1 M HCL, for the same final concentration of 3% DMSO and 3.5 mM HCL as above. After allocation of mice into their respective cages on the day of birth, measurements were performed blindly. The experiment was started with 17 WT animals receiving vehicle (DMSO), 8 WT animals receiving infigratinib, 7 mutant animals receiving vehicle, and 6 mutant animals receiving infigratinib.

### Statistics

Statistical analysis was performed using GraphPad Prism version 10 (GraphPad Software Inc.). Differences between two groups were tested using a two-tailed Student’s t-test for comparing parametric variables. P values < 0.05 were considered significant. Data are presented as the mean ± SD. For the preclinical infigratinib in vivo study, a mixed model for repeated measures with time x treatment interaction effect was used.

### Study approval

Mice were maintained in a specific pathogen-free AAALAC-accredited facility, and all procedures were approved by the Institutional Animal Care and Use Committee (ACUC) of the National Human Genome Research Institute (NHGRI) or the National Institute of Dental and Craniofacial Research (NIDCR).

### Data availability

All data generated or analyzed in this study are included in the manuscript and supporting files. A Supporting Data Values file has been provided for all numerical data. RNA-Seq data were deposited in the NCBI’s Gene Expression Omnibus (GEO) database (GEO GSE306640).

## Author contributions

GA and CRF conceptualized the study. GA, RK, GE, DWC, LG, LFdC and CRF designed research studies. GA, RK, RG, GE, LFdC and CRF conducted experiments. GA, RK, CR, IG, SC, RG, AM, LL, LFdC and CRF acquired data. GA, RK, MR, AC, LFdC and CRF analyzed data. GA, RK, IH, LFdC and CRF wrote the original draft of the manuscript. All authors reviewed and edited the manuscript.

## Acknowledgments

This work was supported by the Intramural Research Program of the National Institutes of Health (NIH) (HD009024 to C.R.F.). The contributions of the NIH authors are considered Works of the United States Government. The findings and conclusions presented in this paper are those of the authors and do not necessarily reflect the views of the NIH or the U.S. Department of Health and Human Services.

