## Supplemental figures for "Disruption of the FGFR1-FGF23-Phosphate Axis and Targeted Therapy in a Murine Model of Osteoglophonic Dysplasia"

Supplemental material:

Supplemental Figure S1 to Supplemental Figure S3;

Supplemental Table ST1, Supplemental Table ST2

Supplemental Data SD1

**Supplemental Figure S1**

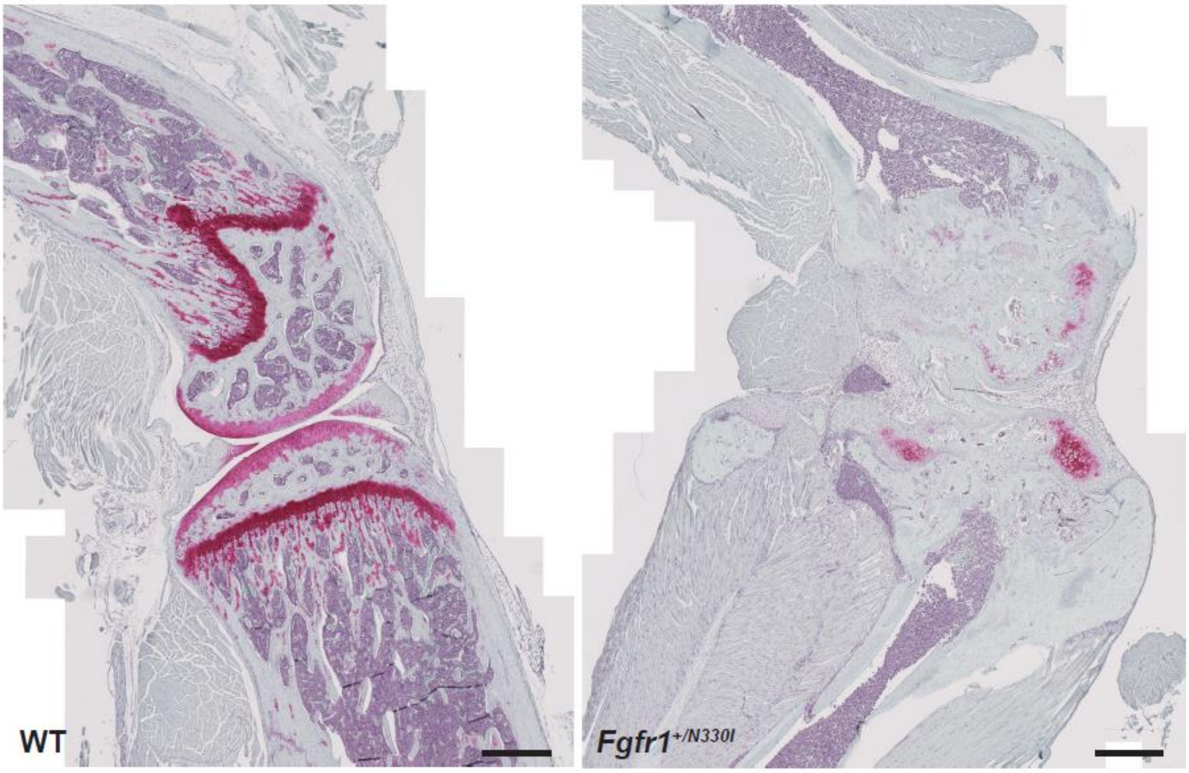


**Supplemental Figure S1. *Fgfr1$^{+/N330I}$* mice display disorganization of cartilage extracellular matrix at 12 weeks.**

Safranin O/Fast green (SafO/FG) staining for proteoglycans showed disorganization of cartilage in 12-week-old *Fgfr1$^{+/N330I}$* mice as indicated by the impaired distribution of the extracellular matrix (red). Scale bar = 500 µm.

### Supplemental Figure S2

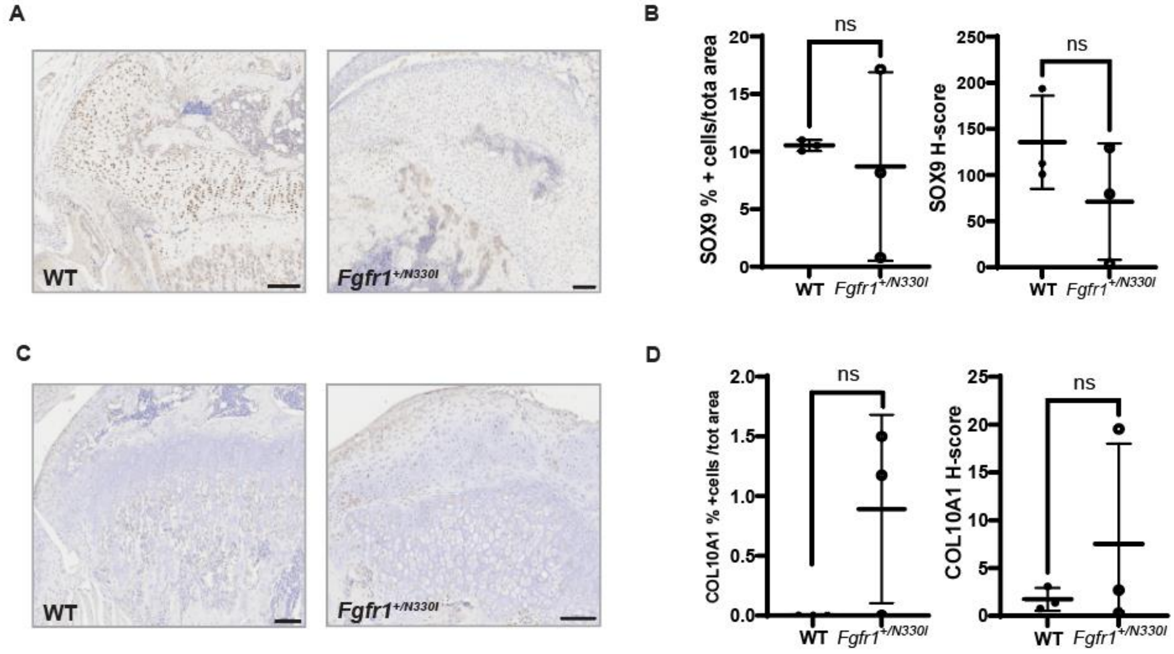


**Supplemental Figure S2. Immunohistochemistry of SOX9 and Collagen X.**

**(A)** Representative photomicrographs and **(B)** staining quantification showing similar protein levels of SOX9 in femurs derived from WT and *Fgfr1*$^{+/N330I}$ mice as shown by quantification of SOX9$^{+}$ cells and relative H-score. **(C)** Representative photomicrographs and **(D)** staining quantification showing similar protein levels of Collagen X (COL10A1) in femurs of WT and *Fgfr1*$^{+/N330I}$ mice. Horizontal and vertical lines represent the mean ± SD of 3 mice/group (ns = not significant). Scale bar = 100 µm.

**Supplemental Figure S3**

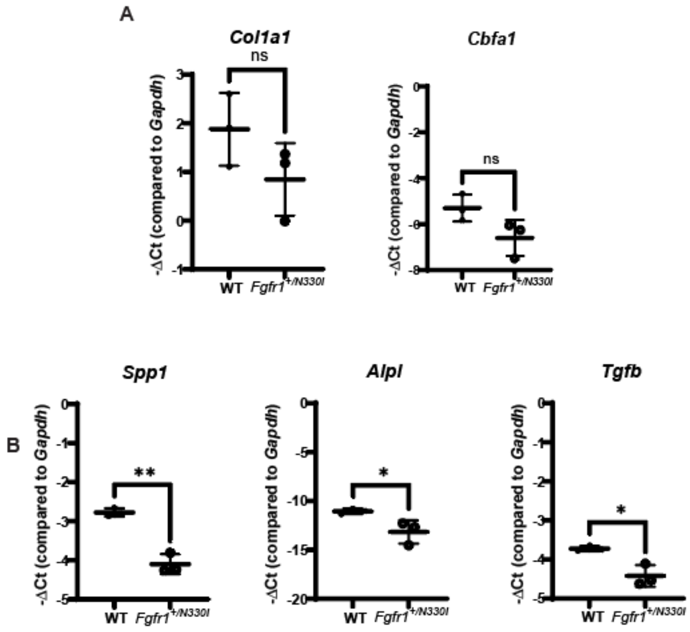


**Supplemental Figure S3.** ***Fgfr1*** $^{+/N330I}$ **mice have reduced mRNA levels of osteoblast differentiation markers.**

**(A)** WT and *Fgfr1* $^{+/N330I}$ mice show comparable messenger RNA (mRNA) expression of Collagen Type 1 (*Col1a1*) and Core-binding factor subunit alpha-1 (*Cbfa1*). **(B)** However, the mRNA levels of important osteogenic markers such as secreted phosphorylated protein 1 (*Spp1*), alkaline phosphatase (*Alpl*) and transforming growth factor beta (*Tgfb*) were significantly downregulated in the mutant mice. Horizontal and vertical lines represent the mean ± SD of 3 mice/group. (ns = not significant, * $p < 0.05$, ** $p < 0.01$).

**Supplemental Table ST1**

| Gene name | Forward sequence | Reverse sequence |
|---|---|---|
| *mCol1a1* | 5'- TGACTGGAAGAGCGGAGAGTA-3' | 5'-AGACGGCTGAGTAGGGAACA-3' |
| *mCbfa1/Runx2* | 5'-AGCCAGGTTCAACGATCTGA-3' | 5'-GGACCGTCCACTGTCACTTT-3' |
| *mSsp1* | 5'-GGCTGAATTCTGAGGGACTAACT-3' | 5'-GCAATGCCAAACAGGCAAAAG-3' |
| *mAlp1* | 5'-CAGGCCGCCTTCATAAGCA-3' | 5'-CTCGTGGGGAGGGTCTGTTC-3' |
| *mTgfb1* | 5'-ATTCCTGGCGTTACCTTGGT-3' | 5'-GTGAGCGCTGAATCGAAAGC-3' |
| *mGapdh* | 5'-GGCAAATTCAACGGCACA-3' | 5'-GTTAGTGGGGTCTCGCTCCTG-3' |

**Supplemental Table ST1.** Primers used for assessing gene expression in mouse cortical bone explants.

**Supplemental Table ST2**

| MicroCT parameter | ***Fgfr1***$^{+/N330I}$ **ossicles** | WT ossicles | p-value |
|---|---|---|---|
| Total volume (mm3) | 31.6 | 13.1 | 0.004 |
| BV/TV (%) | 53.3 | 7.9 | <0.0001 |
| Tb.N (mm) | 8.3 | 2.4 | <0.0001 |
| Tb.Th (mm) | 0.09 | 0.06 | <0.0001 |
| Tb.Sp (mm) | 0.13 | 0.45 | 0.0004 |
| Conn.D (mm$^3$) | 342.0 | 67.6 | 0.0001 |
| BS/BV (mm) | 23.2 | 51.0 | p<0.0001 |
| TMD (mgHA/mm$^3$) | 690.8 | 772.3 | 0.002 |
| DA (dimensionless) | 1.09 | 1.32 | 0.001 |

Abbreviations: BV/TV, Bone volume/total volume; BS/BV, Bone surface/bone volume; Conn.D, Connectivity density; DA, Degree of anisotropy; TMD, Total mineral density; Tb.N, Trabecular number; Tb.Sp, Trabecular spacing; Tb.Th, Trabecular thickness.

**Supplemental Table ST2.** Microarchitectural parameters of in vivo ectopic ossicles (n = 6/cohort).

**Supplemental Data SD1**

AAACCCAGGAACTCCCCCTGCATGGCCCTCAAAGCCAGAGCAGAAAAGGGTACGCAGC
AGACGGGAGCCGTCCCTTTCTGCCTCTCCCCAGCCCTGTAACGGATGCGCTGCTCCCA
TGGCAAAAGGCAGGACCCGGAAAGGACCAGTGCACTGTATACCTTCCAGAACGGTCAA
CCATGCAGAGTGATGGGAGAGTCCGATAGAGTTACCCGCCAAGCACGTATACTCCCCCG
CATCCTCAAAGGAGACAATGCGTAGATGAAGCACCTCCATTTCCTTGTCGGTGGTATTAA
CTCCAGCAGTCTAGAAGAGACGACGGAAGCAAAATGGACAAGCCCGGGACATGAGACC
TCCGAGAGGCGGAGAAACGGAGGGGACAGAGGAGGGGACCAAAACCAGAAGAAGAAA
CGCTCCCCAGCATCTCCTTTGCAACAAGGCAAAAAAACCACACCACCAGCCAGCGAGA
CCTGTTAGAGGAAAACAGCGAGGCTCTGGAGGGGA

**Supplemental Data SD1:** Sequence of the homology-directed repair template used for generation of *Fgfr1*$^{+/N330I}$ mice.
